# GATA2 deficiency during embryogenesis elevates Interferon Regulatory Factor-8 to subvert a progenitor cell differentiation program

**DOI:** 10.1101/2021.01.29.428803

**Authors:** Kirby D. Johnson, Emery H. Bresnick

## Abstract

Cell type-specific transcription factors control stem and progenitor cell transitions by establishing networks containing hundreds of genes and proteins. Network complexity renders it challenging to discover essential versus modulatory or redundant components. This scenario is exemplified by GATA2 regulation of hematopoiesis during embryogenesis. Previously, we demonstrated that loss of *Gata2*, −77 enhancer disrupts the GATA2-dependent genetic network governing erythro-myeloid differentiation. The aberrant network includes the transcription factor Interferon Regulatory Factor-8 and a host of innate immune regulators. Mutant progenitors lose the capacity to balance production of diverse myelo-erythroid progeny. To elucidate mechanisms, we asked if IRF8 is essential, contributory or not required. *Irf8* ablation, in the context of the −77 mutant allele, reversed granulocytic deficiencies of −77^−/−^ embryos and rescued an imbalance of dendritic cell progenitors. Despite many dysregulated components that control vital processes, including transcription and signaling, aberrant elevation of a single transcription factor deconstructed the differentiation program.

## Introduction

The cloning of sequence-specific DNA binding proteins unveiled “master regulators” including GATA1, which promotes erythrocyte, megakaryocyte, mast cell and basophil development, and GATA2, which mediates hematopoietic stem and progenitor cell (HSPC) genesis/function (1). During mouse embryogenesis, GATA2 ablation disrupts multi-lineage hematopoiesis, yielding lethality (2). Human *GATA2* heterozygous mutations cause GATA2 deficiency syndromes involving cytopenias, immunodeficiency, bone marrow failure, leukemia and lymphedema (3). These mutations alter the *GATA2* coding region or “+9.5” intronic enhancer (4). Another pathogenic enhancer resides upstream of *GATA2* (77 kb in mice; 110 kb in humans) (5). A 3q21;q26 inversion removes this enhancer from one allele, positioning it near *MECOM1*, inducing EVI1 expression and AML (6,7). Homozygous deletion of the −77 enhancer depletes megakaryocyte-erythroid progenitors (MEPs), yielding embryonic lethality (5,8). −77^−/−^ myelo-erythroid progenitors lose multi-lineage differentiation and generate predominantly macrophages *ex vivo* (5). Monocytopenia characterizes GATA2 deficiency syndrome, but bone marrow macrophages persist (9).

Although GATA2-regulated genes have been described (1), downstream mediators and epistatic relationships that control HSPCs are unresolved. Discovering mediators has been challenging due to context-dependent GATA2 mechanisms (1). Proteomics and single-cell transcriptomics with −77^−/−^ progenitors revealed losses of proteins mediating erythroid, megakaryocyte, basophil and granulocyte differentiation and elevated innate immune proteins, including Interferon Responsive Factor 8 (IRF8) (10).

IRF8 interacts with other IRFs, PU.1, AP-1 and BatF3 at composite binding sites to control immune cell development (11). IRF8 levels dictate target gene selection by regulating transcription factor complex composition (11). As IRF8 promotes monocyte development, and its loss favors neutrophil generation (12,13), it is attractive to propose that IRF8 upregulation in −77^−/−^ progenitors unrestrains monocytic differentiation, and GATA2 downregulation of IRF8 is critical for multi-lineage differentiation. Since expression correlations often do not reflect causation, and IRF8 function in GATA2 networks is not understood, IRF8 might function non-redundantly or redundantly. We asked if *Irf8* ablation, in the context of the −77^−/−^ allele, rescues the diverse myelo-erythoid differentiation potential of progenitors.

## Results and Discussion

The murine granulocyte-monocyte progenitor (GMP) population (Lin^−^cKit^+^Sca-1^−^CD34^+^FcγR(CD16/CD32)^hi^) contains bipotential hematopoietic progenitors and committed granulocyte (GP) and monocyte (MP/cMoP) progenitors (14). In comparison to wild type fetal liver, the GP:MP ratio within the GMP population of −77^−/−^ fetal liver favors MPs (10), and −77^−/−^ progenitors generate predominantly macrophages *ex vivo* (10). IRF8 regulates survival and differentiation, but not production, of lineage-committed bone marrow progenitors (14). IRF8 deficiency increases granulocytes, whereas IRF8 promotes monocyte generation (14).

To determine if high IRF8 resulting from −77 enhancer loss causes the GP:MP imbalance or contributed to deficiencies in other progenitor populations, fetal liver progenitors were quantified in −77 and *Irf8* double-mutant (−77^−/−^;*Irf8*^−/−^) embryos (Figure 1A). In red cell-depleted E14.5 fetal liver, *Irf8* expression was undetectable in −77^+/+^;*Irf8*^−/−^ and −77^−/−^;*Irf8*^−/−^ embryos and elevated in −77^−/−^;*Irf8*^+/+^ versus −77^+/+^;*Irf8*^+/+^ littermates (Figure 1B). As *Gata2* expression was indistinguishable in −77^−/−^;*Irf8*^+/+^ and −77^−/−^;*Irf8*^−/−^ livers, changes in progenitor populations in double mutants were not caused by increased *Gata2* expression. IRF8 loss did not rescue the −77^−/−^ megakaryocyte-erythroid progenitor (MEP) deficiency (7) (Figure 2A, B). GPs (CD115^low^) and MPs (CD115^hi^) are abundant within the wild type fetal liver GMP population (10), and −77 enhancer deletion shifted the balance to favor MPs (Figure 2A, C). IRF8 loss greatly reduced the percentage of MPs and elevated GPs within the Ly6C^−^ GMP pool. As a percentage of the Lin^−^ population, −77^−/−^; *Irf8*^−/−^ MPs were reduced by 29% compared to 77^−/−^;*Irf8*^+/+^ littermates. GPs increased 9.3 fold to comprise 20% of the Lin^−^ pool (Fig 2D). As GPs dominated in −77^−/−^;*Irf8^−/−^* mutants, reducing IRF8 reversed the imbalance in myeloid progenitors that favored MPs in 77^−/−^ embryos.

**Figure 1.**
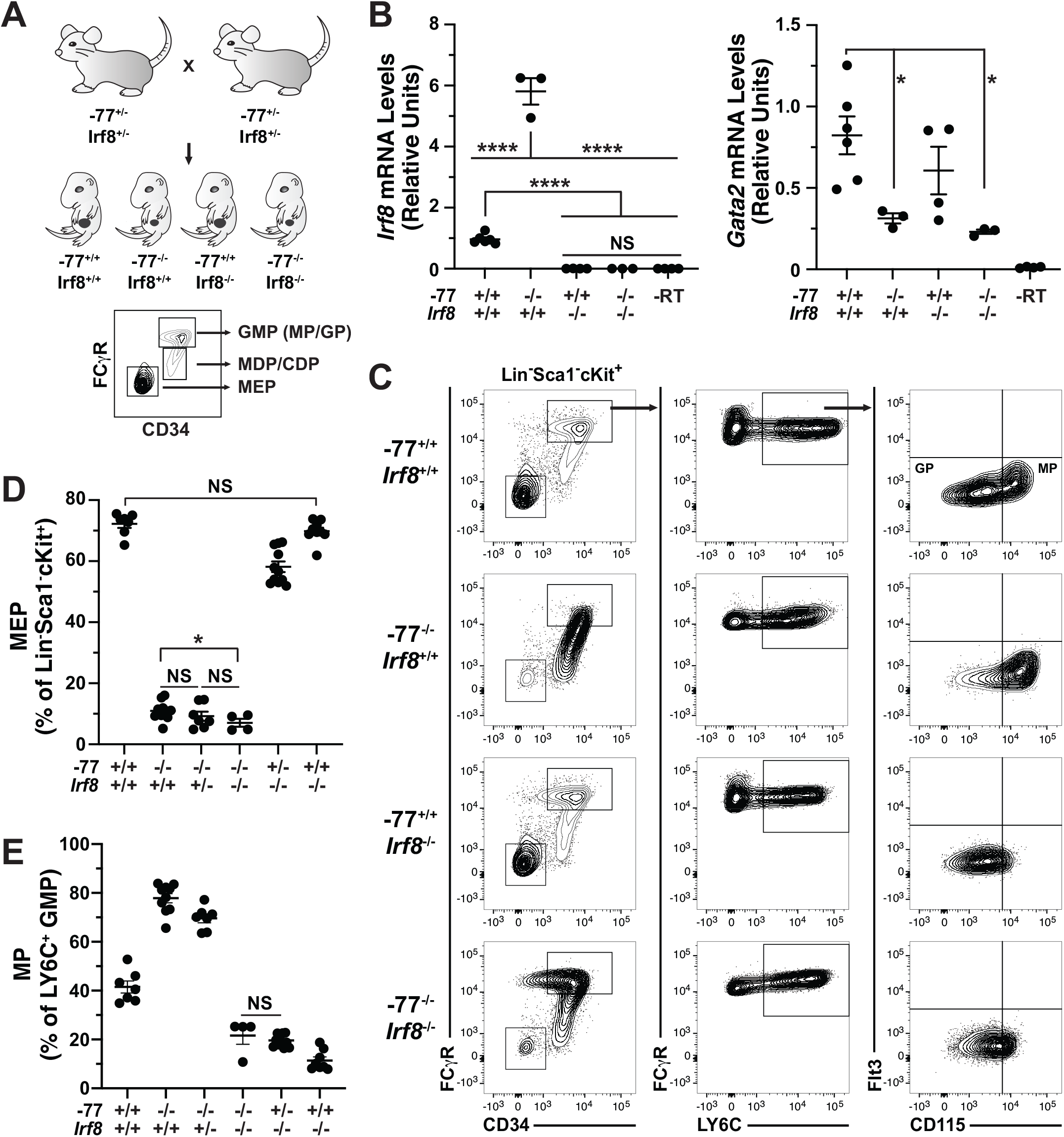
Double mutant *in vivo* rescue system. (A) Experimental strategy. Mice heterozygous for −77 enhancer deletion or *Irf8^tm1.2Hm^* were bred to generate E14.5 embryos. Fetal liver progenitor populations with select genotypes were analyzed by flow cytometry. Error bars represent mean ± SEM (3-11 embryos; 7 litters). Statistics were calculated using unpaired two-tailed Student’s t test. ****, P < 0.0001; *, P < 0.05. NS, not significant (P ≥ 0.05). (B) RT-qPCR analysis of *Irf8* and *Gata2* expression in red cell-depleted fetal livers. *Irf8* primers (GGCAAGCAGGATTACAATCAG and CCACACTCCATCTCAGGAAC) detected exon loss in *Irf8*^−/−^ embryos. *Gata2* primers were described previously (5). P < 0.01 unless indicated.

**Figure 2.**
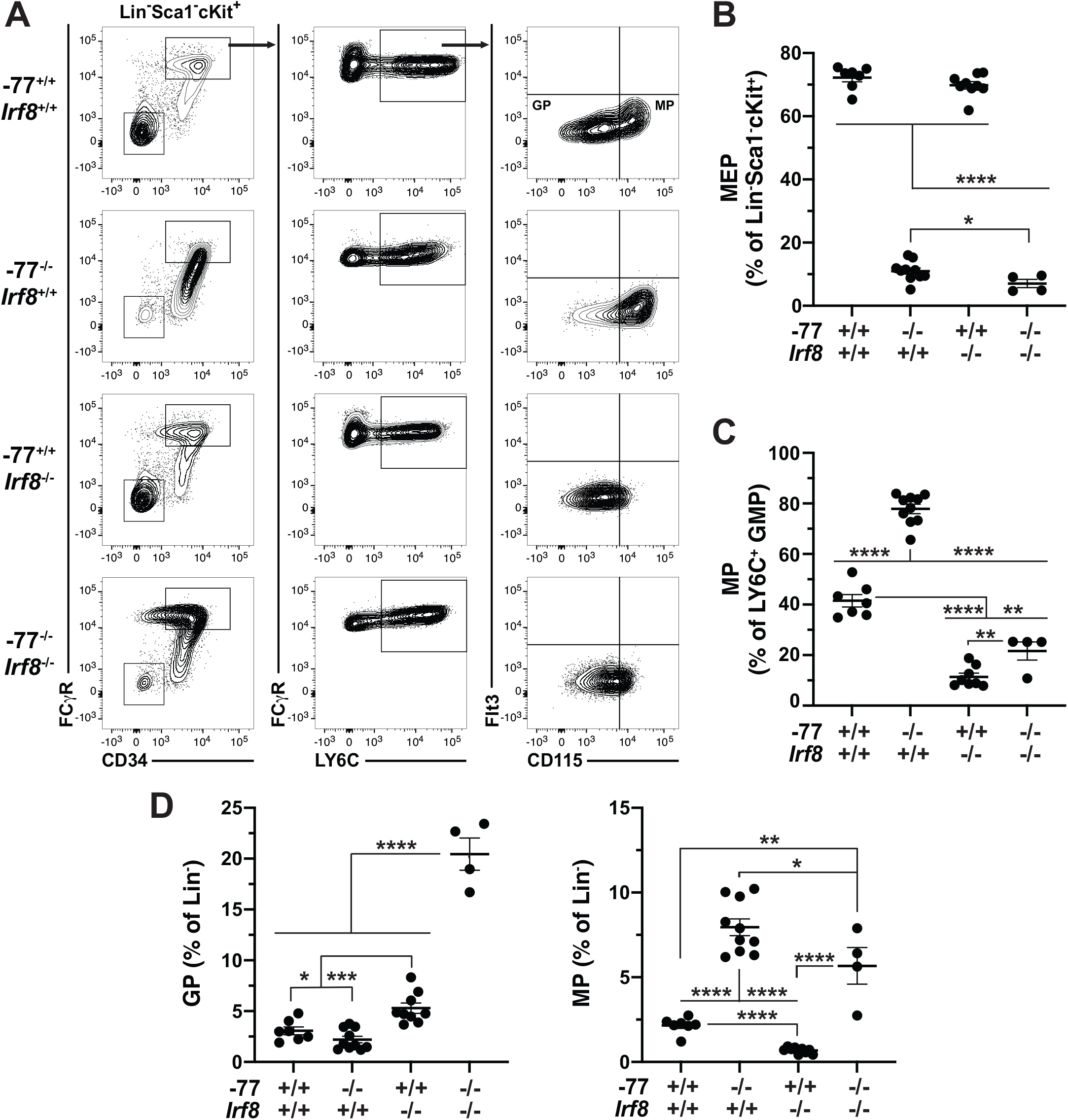
Elevated Interferon Regulatory Protein-8 causes aberrant myeloid differentiation potential of GATA2-deficient progenitor cells. (A) Representative flow cytometry analysis (MEP, GP, and MP populations). GPs and MPs are distinguished from bipotential GMPs by Ly6C and from each other by M-CSF receptor (CD115). Quantification of the frequency of MEP (B), MP (C and D) and GP (D) progenitor populations from 4-11 embryos obtained from 7 litters. Error bars represent mean ± SEM. Statistics were calculated using unpaired two-tailed Student’s t test. *, P < 0.05; **, P < 0.01; ***, P < 0.001; ****, P < 0.0001.

Dendritic cell (DC) defects characterize GATA2 deficiency syndrome (3). As with monocytes, IRF8 controls DC generation (14). IRF8 levels within DC-restricted progenitors (CDPs) determine the DC subtype produced; high and low IRF8 promotes cDC1 and cDC2 production, respectively (11). *Irf8*^−/−^ mice exhibit increased bone marrow monocyte-dendritic cell progenitors (MDPs) and reduced CDPs (14).

In addition to elevated levels of innate immune and monocyte genes (10), transcriptomic analysis of −77^−/−^ fetal liver progenitors revealed increased expression of multiple genes important for DC differentiation and/or selectively enriched in DC progenitor or precursor cells (Figure 3A) (15). Since IRF8 promotes DC development in adult mice and humans (14,16), GATA2 suppresses IRF8 expression in mouse embryos and high IRF8 corrupts GATA2-deficient (−77^−/−^) progenitor differentiation, we asked if the GATA2-IRF8 axis impacts DC progenitors. MDPs (cKit^hi^) produce CDPs, which express intermediate levels of cKit (17). In *Irf8*^−/−^ E14.5 fetal liver, CDPs and MDPs comprised 9.3% and 91%, respectively, of the FcγR^low^ CD34^+^ Flt3^+^ CD115^+^ Ly6C^−^ progenitors (Figure 3B, C). In wild type littermates, CDPs were 21% of the total, contrasting with 55% in −77^−/−^ fetal liver (p = 5.8^−7^). Bi-allelic *Irf8* and −77 loss restored the CDP:MDP ratio to resemble wild type embryos. However, as with MPs, the proportion of MDPs and CDPs in the 77^−/−^ Lin^−^ pool increased; 4.8 fold (P =1.5E^−5^) and 21 fold (P = 1.2E^−6^), respectively, compared to 77^+/+^ (Figure 3D). In −77; *Irf8* double mutants, MDPs were reduced by 44% (P = 0.03), while the percentage of CDPs decreased 7.5 fold (P = 0.0002), consistent with the role of IRF8 in CDP production (12). Opposing GATA2 and IRF8 activities controlled the CDP:MDP ratio, extending GATA2-IRF8 axis function beyond establishing a balance between MPs and GPs.

**Figure 3.**
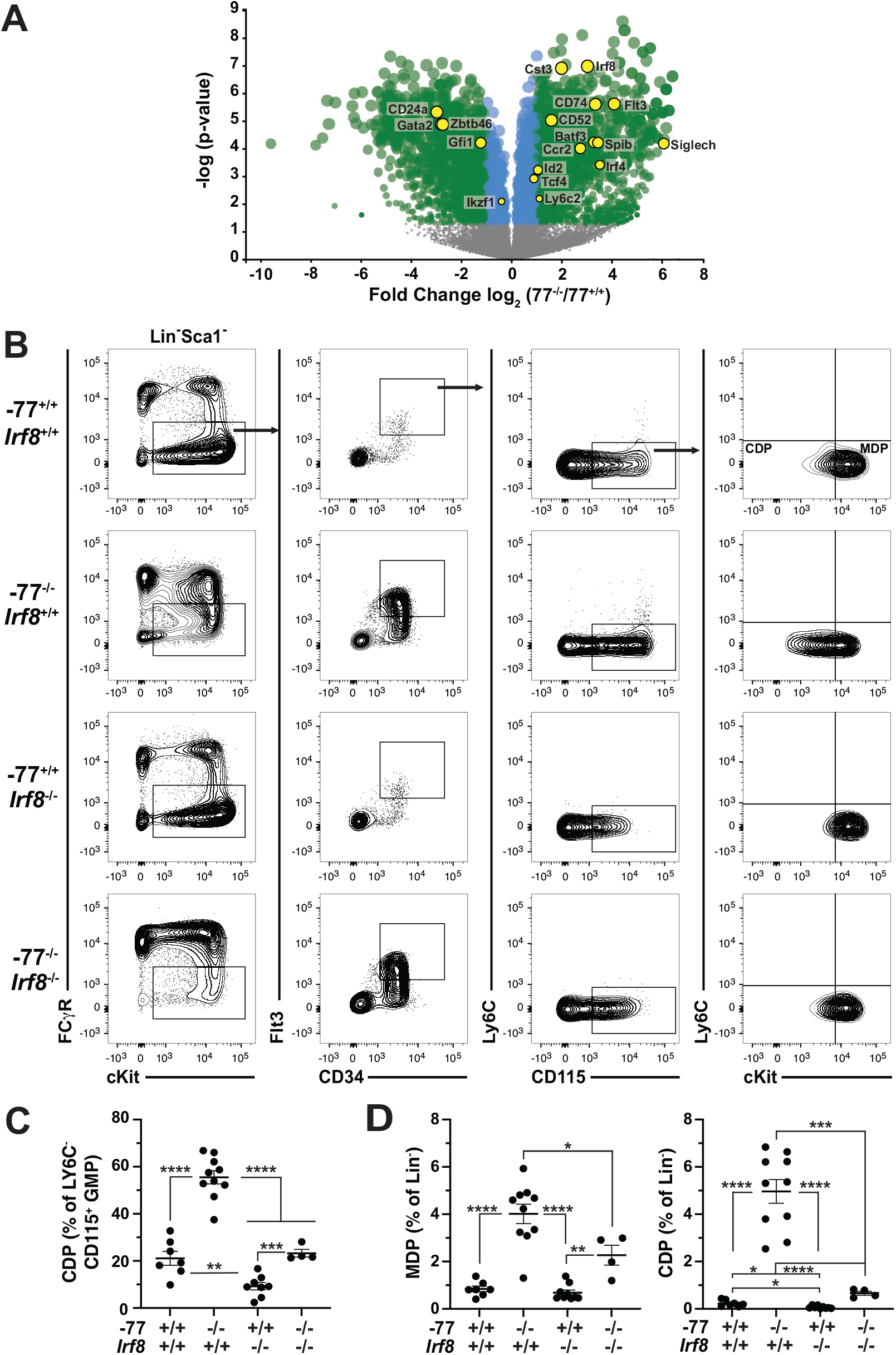
GATA2 suppression of Interferon Regulatory Factor-8 levels restricts common dendritic cell progenitor production. (A) Volcano plot of differentially expressed genes from transcriptome analysis of lineage-depleted wild type and −77^−/−^ fetal livers cultured for 3 d (10). GEO accession: GSE133606. DC genes included factors involved in DC differentiation and/or differentially expressed in DC progenitor populations (15). *Gata2* downregulation, resulting from its −77 enhancer deletion, is also depicted. (B) Representative flow cytometry analysis of monocyte-dendritic cell progenitor (MDP) and common dendritic cell progenitor (CDP) populations. Quantitation of the frequency of CDP (C and D), and MDP (D) progenitor populations from 4-11 embryos obtained from 7 litters. Error bars represent mean ± SEM. Statistics were calculated using unpaired two-tailed Student’s t test. P values were less than 0.01 except where indicated. *, P < 0.05; **, P < 0.01; ***, P < 0.001; ****, P < 0.0001.

Our results provide evidence for a GATA2-IRF8 axis that dictates myelo-erythroid progenitor generation and function during mouse embryogenesis. Since *Gata2* and *Irf8* mRNAs co-reside in single progenitors at reciprocal levels, and re-introducing GATA2 into −77^−/−^ progenitors decreases IRF8 (10), the levels of GATA2 and IRF8 in a single cell are key determinants of this developmental mechanism. During basophil development, IRF8 loss blocks Lin^−^Sca−1^−^cKit^+^CD150^−^β7integrin^−^CD27^+^ GP differentiation, and GATA2 expression in *Irf8*^−/−^ bone marrow cells rescues basophil differentiation (18). Whether GATA2 and IRF8 function in the same or distinct networks and cells to control differentiation in this context was not described.

The reduced GATA2 of −77^−/−^ myelo-erythroid progenitors, which impedes erythroid, megakaryocyte, granulocyte and basophil developmental trajectories, while maintaining monocyte progenitors, is associated with collapse of the GATA2 network (3,161 differentially expressed genes and 434 differentially expressed proteins in −77^−/−^ versus −77^+/+^ progenitors) (10). As the aberrant network includes many regulatory factors, one would assume that a multi-component mechanism skews differentiation. Our results with the double-knockout rescue paradigm demonstrated that ablating *Irf8* rectified the GP deficiency in −77^−/−^ progenitors (Figure 4), thus establishing a mechanism in which GATA2 suppresses IRF8 levels to control myelo-erythroid progenitor fate, and IRF8 elevation in GATA2-deficient embryos corrupts this mechanism. Applying the strategy innovated to other network components will discriminate vital stem and progenitor cell regulators from those exhibiting intriguing expression patterns that represent inconsequential correlations.

**Figure 4.**
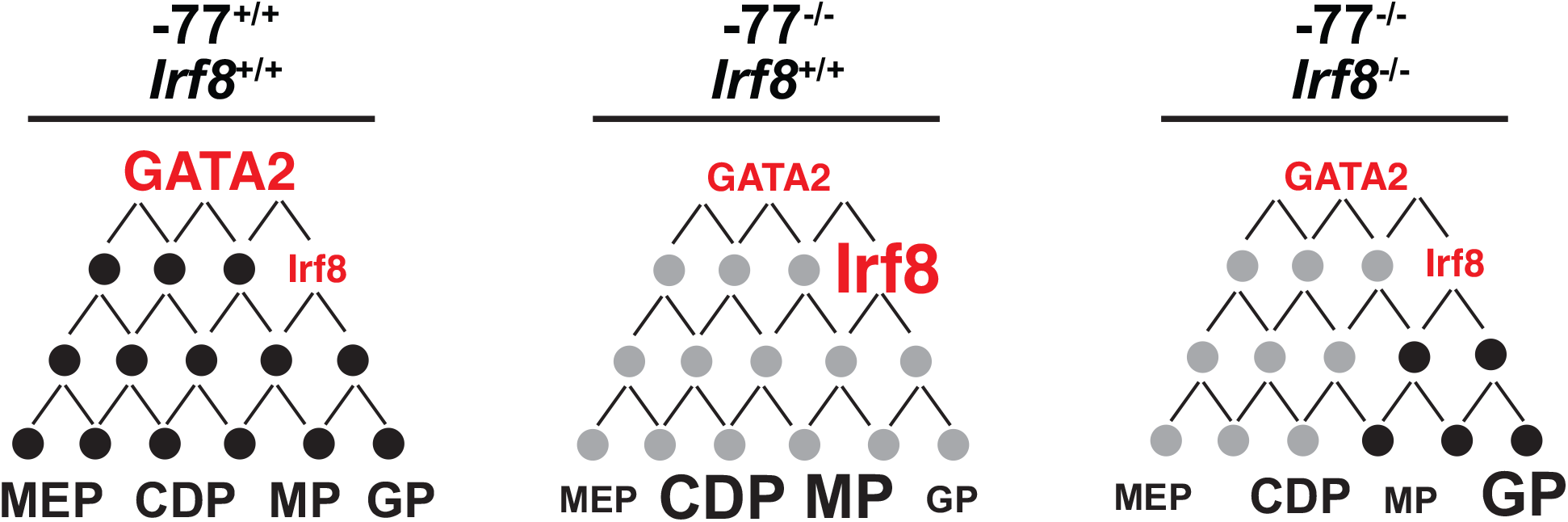
Model illustrating the function of Interferon Regulatory Factor-8 as an essential mediator within GATA2 genetic networks. Physiological levels of GATA2 establish gene regulatory networks (black circles) with low *Irf8* expression, which maintains a balance in myeloid and dendritic cell progenitors. Loss of the −77 enhancer and consequent reduction in GATA2 disrupts the network (gray circles), upregulating *Irf8* and increasing proportions of MP, MDP and CDP populations. Loss of *Irf8* in the context of −77^−/−^ rescued the macrophage and dendritic progenitor overproduction.

## Materials and Methods

### Mice

*Gata2* −77^−/−^ mice (10) and *Irf8*^−/−^ strain B6(Cg)-*Irf8^tm1.2Hm^*/J (Jackson Labs, Bar Harbor, Maine) were bred to generate −77^+/−^;*Irf8*^+/−^ males and females for timed matings. Animal protocols were approved by the UW–Madison Institutional Animal Care and Use Committee in accordance with the Association for Assessment and Accreditation of Laboratory Animal Care (AAALAC International) regulations.

### Flow cytometry

E14.5 fetal liver MEPs, MPs, GPs, MDPs and CDPs were quantified using the LSR Fortessa flow cytometer (BD Biosciences) or sorted using a FACSAria (BD Biosciences). Antibodies were from Biolegend unless indicated. B220, TER-119, CD5, CD11b, IgM, Ly6G and Sca1 antibodies were FITC-conjugated. Other antibodies were Blue Violet (BV) 605-conjugated CD16/CD32 (BD Biosciences), BV711-conjugated Ly-6C, phycoerythrin (PE)-conjugated CD115, eFluor 660–conjugated CD34 (Thermo Fisher Scientific), peridinin chlorophyll (PerCP)-efluor710–conjugated CD135 (Thermo Fisher Scientific) and PE-Cy7–conjugated cKit. Stained cells were washed with PBS, 2% FBS, 10 mM glucose, and 2.5 mM EDTA, resuspended in the same buffer containing DAPI and passed through 25 μm strainers. DAPI (4′,6-diamidino-2-phenylindole) was used for live-dead discrimination.

## Acknowledgments

NIH grant R01DK68634, Carbone Cancer Center P30CA014520 and Edward Evans Foundation grants supported this work. We thank Mabel M. Jung for graphic design.

## Competing Interests

The authors have nothing to declare.

## Notes

### Competing Interest Statement

The authors have declared no competing interest.

